# Exploring Perceptual Illusions in Deep Neural Networks

**DOI:** 10.1101/687905

**Authors:** Emily J. Ward

**Affiliations:** Department of Psychology, University of Wisconsin - Madison, 1202 W. Johnson Street, Madison, WI 53703 USA

**Keywords:** perception, illusions, deep neural networks

## Abstract

Perceptual illusions—discrepancies between what exists externally and what we actually see—reveal a great deal about how the perceptual system functions. Rather than failures of perception, illusions expose automatic computations and biases in visual processing that help make better decisions from visual information to achieve our perceptual goals. Recognizing objects is one such perceptual goal that is shared between humans and certain Deep Convolutional Neural Networks, which can reach human-level performance. Do neural networks trained exclusively for object recognition “perceive” visual illusions, simply as a result of solving this one perceptual problem? Here, I showed four classic illusions to humans and a pre-trained neural network to see if the network exhibits similar perceptual biases. I found that deep neural networks trained exclusively for object recognition exhibit the Müller-Lyer illusion, but not other illusions. This result shows that some perceptual computations that are similar to humans’ may come “for free” in a system with perceptual goals similar to humans’.

## Introduction

Why do we see perceptual illusions? Rather than random failures, illusions are systematic and persistent distortions that cause us to see the world in a way that differs from how it actually is. Do illusions arise as a result of solving one specific type of perceptual problem, or are illusions a consequence of computations necessary in a system that has many perceptual goals and must solve many types of problems?

Certain types of Deep Convolutional Neural Networks have been built and trained to solve a critical and computationally difficult problem of recognizing many kinds of objects, and therefore share one perceptual goal with humans. At this task, these networks have reached or exceeded human performance (e.g. Simonyan and Zisserman (2014); Szegedy, Vanhoucke, Ioffe, Shlens, and Wojna (2015)). However, deep convolutional neural networks only approximate human neurophysiology (both these networks and the human brain are grossly hierarchical) and poorly approximate neural information prorogation (there is no back propagation in these networks). Thus, given the excellent behavioral performance but poor neurophysiological match, it is unknown how similar the representations and computations are between deep neural networks and the human brain.

It is possible that a perceptual goal, such as object recognition, is critical enough to cause similar representations and computations to emerge in any system that shares that goal. For example, deep convolutional neural networks develop selectivity for many of the same features important in early and higher-level visual processing (Cadena et al., 2019; Yamins et al., 2014). Does the similarity in selectivity between these networks and humans suggest similarity in perceptual computation? Specifically, do neural networks trained exclusively on object recognition develop that perceptual biases that would cause them to “perceive” visual illusions, or other special cases in perception?

My approach was to compare the strength of classic illusions (Figure 1) in humans and in a neural network (VGG19, Simonyan and Zisserman (2014)) pre-trained exclusively on ImageNet object classification. I assessed human and network perception of the illusions using similar paradigms for both types of “agents”: human participants viewed an illusion (such as a Müller-Lyer illusion, with arrows pointing out) and chose an item that best matched the relevant feature of the illusion (such as choosing which horizontal line best matched the horizontal line in the illusion); the neural network processed the same image of the illusion and the items, and the best match was determined by how similar the test item was to the illusion at the final layer of the network (given by cosine distance between the two associated feature vectors).

**Figure 1:**
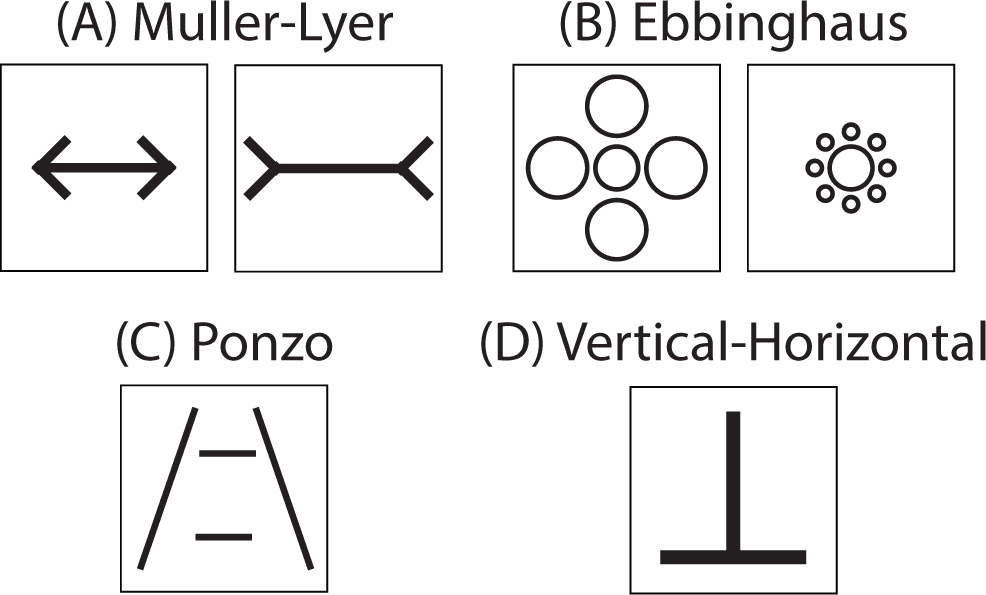
The illusions tested were for classic illusions: the Müller-Lyer (A), Ebbinghaus (B), Ponzo (C), and Vertical-Horizontal Illusion (D).

Overall, I found that the network exhibited the Müller-Lyer illusion, but not other illusions.

## Methods

### Participants

#### Humans

Observers were recruited and run online via Amazon Mechanical-Turk. Observers who failed one of two catch trials were excluded from the analyses. Overall, 16% of people were excluded.

#### Neural Network

The primary neural network used for this analysis was VGG19 (Simonyan & Zisserman, 2014), constructed in the Keras python package (Chollet et al., 2015). The network has 19 layers typically classifies objects into 1000 object categories. For my analyses, I extracted the feature vector from the last fully connected layer (FC7), before labels were applied. For comparison, the analyses were also run on VGG16 and InceptionV3 (Szegedy et al., 2015), both also pre-trained on ImageNet (Deng et al., 2009).

### Stimuli

Four geometric illusions (Müller-Lyer, Ebbinghaus, Ponzo, and Vertical-horizontal) were used (Figure 1A-D). The images were grayscale and 448 px x 448 px, although they were scaled for different networks. All stimuli were generated in various forms to promote generalization.

#### Illusions

For the Müller-Lyer illusion (Figure 1A), 4 parameters (central line length, arrow length, orientation, and stroke width) of the illusion varied: The total length of the horizontal line was either 150 or 200 px. The arrows had a length of 20, 30 or 40 px. The illusion was either horizontal or vertical. The stroke width of the lines was either 3, 6, or 9 px.

For the Ebbinghaus illusion (Figure 1B), 3 parameters (central circle radius, surrounding circle radius, and stroke width) of the illusion varied: The radius of the target circle was either 20, 30, or 35 px. The radius of the smaller inducers was 7 or 20 px and the radius of the larger inducers was 15 or 50 px. The stroke width of the lines was either 2, 3, or 4 px.

For the Ponzo illusion (Figure 1C), 2 parameters (bounding line slope and stroke width) of the illusion varied as follows: the slope of the two bounding lines was 20, 28, or 37. The two horizontal lines were always 80 px. The stroke width of the lines was either 4, 6, or 8 px.

For the Vertical-horizontal illusion (Figure 1D), 2 parameters (orientation and stroke width) of the illusion varied as follows: the illusion could either be presented upright or upside down and the stroke width of the lines was either 4, 6, 8, or 10 px. The length of the horizontal and vertical liens were always 200 px.

### Procedure

#### Humans

On each trial, participants viewed an illusion and chose one of two items that best matched the relevant feature of the illusion. The illusion was presented at the top center of the browser window and the two items were presented below it in a row. Participants were instructed to pay attention to the relevant feature of the illusion (e.g. horizontal line) and pick the item that matched it (e.g. in terms of length). Participants indicated their choice by selecting a radio button on the page.

In total each participant completed 25-30 trials (preceded by two easy practice trials and succeeded by two catch trials, identical to the practice). On all trials, the test illusion remained the same (no change to its parameters, such as size, stroke width etc.) and only the test items differed from trial to trial: one item was always an identical match (e.g. in terms of length) and the other was always longer or shorter. Participants saw each length of the test item once per experiment, with order and display side (left or right side of the browser) randomized.

#### Neural Networks

To fairly compare human and network performance, I ran similar trials through the network. The network processed images of the target illusion and the two test items. As with human participants, one item was an identical match and the other was longer or shorter. For each image, I extracted the features from the last fully-connected layer, FC7. I then computed the cosine distance between the identical item and the illusion and the distance between the test item and the illusion. Whichever item produced a smaller distance (i.e. best matched the illusion) was the choice of the network. In total, the network saw all test item lengths, and all combination of illusion parameters (e.g. stroke width, inducer size, etc.).

### Analysis

Each trial generated a binary output (1 = “chose longer item”; 0 = “did not choose longer item”) for both the human participants and and the network. The probability of choosing the test item of a particular length was modeled using logistic mixed effects regression, as test item length and agent (human and network) as predictors and participant as a random effect. The 50% point of the logistic curve was used to estimate the magnitude of the illusion. If, for example, the illusion is seen longer than it is, then the longer item will continue to be chosen beyond what would be expected by chance, if the illusion was seen veridically. This approach allowed me to determine whether neural networks “perceive” visual illusions in any form and whether what they perceive matches what what humans perceive.

## Results

Because my two-item paradigm differs from the typical method for measuring illusions (which is done by adjusting the illusion; e.g. Axelrod, Schwarzkopf, Gilaie-Dotan, and Rees (2017)), I first confirmed the presence of the illusion in humans, then compare it to the network behavior.

I found that both human participants and neural networks exhibited the Müller-Lyer illusion (Figure 2A). Humans judged the outward facing arrows as 16% shorter than veridical and the inward facing arrows as 22% longer than veridical, thus demonstrating the illusion. The network also differentiated between the two versions of the illusion, although it’s response was shifted compared to humans: it judged the outward version as 37% shorter than veridical and the outward version as 5% shorter. Overall, however, network behavior differed from humans’, confirmed by by a significant agent (human and neural network) by test item length interaction (*b*=1.987, *t* =3.693, *p*<0.001).

**Figure 2:**
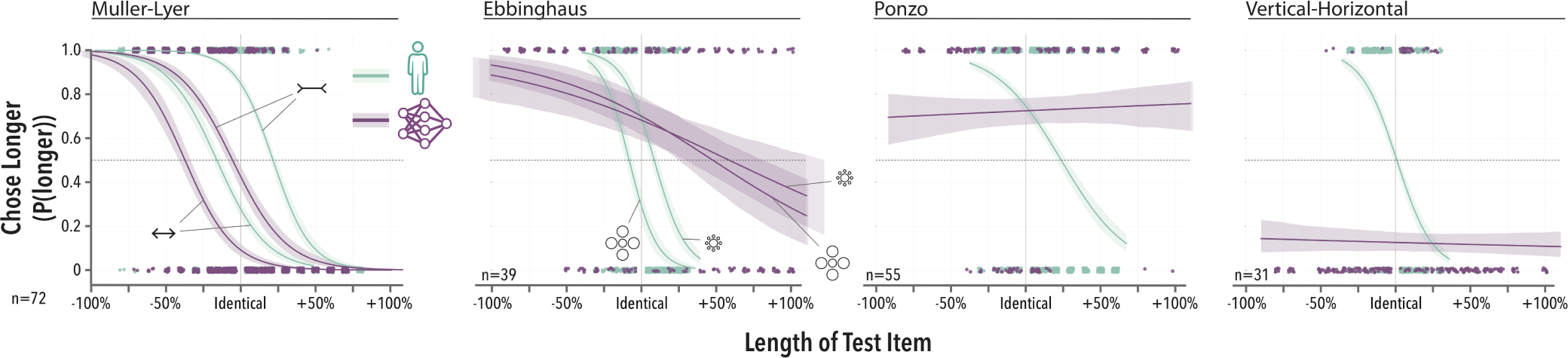
The probability of choosing the longer (or larger) test item as a function of the length of the test item (%). Human (cyan) and neural network (purple) behavior is plotted for four classic illusions: Müller-Lyer, Ebbinghaus, Ponzo, and Vertical-horizontal.

Humans exhibited the Ebbinghaus illusion, but the network only demonstrated some ability to differentiate the two versions of the illusion (Figure 2B). Humans judged the central circle surrounded by large inducers as 8% smaller than veridical and the circle surrounded by smaller inducers as 9% larger than veridical, thus demonstrating the illusion. The network’s response the two versions of the illusion was small and shifted compared to humans: the large-inducers circle was 49% larger than veridical and the smaller-inducers circle was 59% larger than veridical. This difference from humans was confirmed by a significant agent by test item length interaction (*b*=11.626, *t* =12.491, *p*<0.001).

In contrast, although humans participants exhibited the Ponzo illusion, the neural network did not (Figure 2C). Humans judged the top horizontal line as 24% larger than veridical, thus demonstrating the illusion, although the magnitude here was much larger than than previous demonstrations (e.g. Axelrod et al.). In the neural network, the magnitude of the illusion (given by the 50% probability point) could not be determined by the logistic function. The difference between human and network behavior was confirmed by a significant agent by test item length interaction (*b*=5.986, *t* =10.890, *p*<0.001).

Finally, neither human participants nor the neural network demonstrated the Vertical-horizontal illusion using this paradigm (Figure 2D). Humans judged the vertical line as only 1% longer than veridical, which is less than the typical magnitude of this illusion. In comparison, the neural network also failed to demonstrate the illusion and its choices were not well fit using a logistic function. The difference between human and network behavior was confirmed by a significant agent by test item length interaction (*b*=9.538, *t* =10.778, *p*<0.001).

### Other Networks

When I repeated these analyses using two other networks, deeper and more complex networks produced behavior closer to human behavior (Figure 3). Across VGG16, VGG19, and InceptionV3, the overall pattern of results was similar, but the agent by test item length interaction terms decreased for the deeper networks (except for Vertical-Horizontal illusion). In fact, for the Müller-Lyer illusion, InceptionV3 matched human behavior, indicated by an non-significant interaction term, (*b*=0.711, *t* =0.743, *p*=0.457).

**Figure 3:**
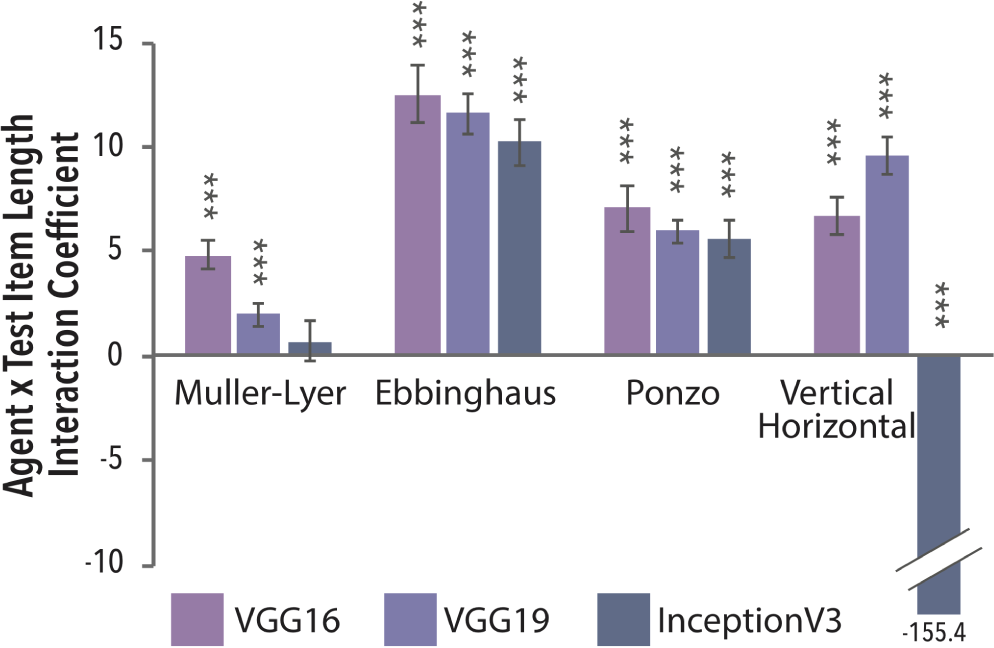
Similarity between human and neural network behavior. The interaction between agent (human and neural network) and test item length is plotted for three neural networks: VGG16 (light purple), VGG19 (purple), and InceptionV3 (dark purple). The smaller the interaction term, the closer match to human behavior.

## Discussion

I found that a deep convolutional neural network (VGG19) trained exclusively for object recognition exhibited the Mϋller-Lyer illusion, despite having had no prior exposure to line drawings of this sort. This finding reveals that neural networks can develop perceptual biases similar to those in human perception, causing them to “perceive” at least one type of illusion.

The Müller-Lyer has previously been demonstrated in other unusual situations, such as in children who have gained sight after extended early-onset blindness (Gandhi, Kali, Ganesh, & Sinha, 2015) and even in blind individuals who feel a haptic version of the illusion (Heller et al., 2002). In contrast, the network did not show similar success with the other classic illusions: while it was able to discriminate the two versions of Ebbinghaus illusion, its behavior was not that similar to human behavior, and for the Ponzo and Vertical-horizontal illusion, the behavior was not comparable to human behavior.

Overall, these results show that neural networks can develop similar perceptual biases as humans by having a shared perceptual goal, such as object recognition, although the extent to which similar computations arise for free is limited.

One limitation with my method is the difficulty in aligning tasks for humans and neural networks. For example, while it is easy to tell human participants to “pay attention to the horizontal line” in a Müller-Lyer illusion, it is difficult to do the same in a neural network that lacks any attentional mechanisms and has not been trained for such a task. There may be evidence of such “misunderstanding” in the results for the Ebbinghaus illusion, where the network judged both illusion types as larger than veridical, possibly due to the fact that the full extend of the display was always larger than the identical test item. However, if it were the case that the network *always* produced responses based on the size of the entire display rather than the feature of interest, then the Müller-Lyer illusion results should also over-estimate the size of the line (because the arrows always extend the total size of the display). That is not the case. Instead, the network systematically under-estimated the size of the lines, for both versions of the illusion. In future, fine-tuning of the network may better ensure that the network is approaching the task properly. However, in this study, the point was specifically to test how much can be gained with the just perceptual goal of object recognition alone, without any additional training.

Beyond showing how neural network perceptual behavior compares with human perceptual behavior, a strength of my approach is that it allows us to isolate the contributions to visual computations made by specific perceptual goals. In this study, I found that the Müller-Lyer illusion can emerge when a system simply needs to recognize many different objects. If I were able to demonstrate the Ponzo illusion in a similar neural network by training it to recognize objects in 3D or by training it to navigate, that would mean that those goals are also critical to human perception – since we irresistibly experience the illusion. In this manner, fine-tuning these network would not only show that neural networks can be made to exhibit the illusions, but it provides new opportunities to identify goals that may otherwise not be giving as much consideration by vision science and cognitive science more generally.

## Acknowledgments

I’d like to acknowledge Gary Lupyan and Chaz Firestone for helpful discussions of this project.

## References

Axelrod, V., Schwarzkopf, D. S., Gilaie-Dotan, S., & Rees, G. (2017, January). Perceptual similarity and the neural correlates of geometrical illusions in human brain structure.Scientific Reports, 7, 39968. Retrieved from https://doi.org/10.1038/srep39968

Cadena, S. A., Denfield, G. H., Walker, E. Y., Gatys, L. A.,Tolias, A. S., Bethge, M., & Ecker, A. S. (2019, April). Deep convolutional models improve predictions of macaqueV1 responses to natural images. PLOS ComputationalBiology, 15(4), e1006897. Retrieved 2019-06-08, from https://journals.plos.org/ploscompbiol/article?id=10.1371 doi:10.1371/journal.pcbi.1006897

Chollet, F., et al. (2015). Keras. https://keras.io.

Deng, J., Dong, W., Socher, R., Li, L.-J., Li, K., & Fei-Fei, L. (2009). ImageNet: A Large-Scale Hierarchical Image Database. In Cvpr09.

Gandhi, T., Kali, A., Ganesh, S., & Sinha, P. (2015, May). Immediate susceptibility to visual illusions after sight onset. Current biology : CB,25(9), R358–R359. Retrieved 2019-06-08, from https://www.ncbi.nlm.nih.gov/pmc/articles/PMC4863640/doi:10.1016/j.cub.2015.03.005

Heller, M. A., Brackett, D. D., Wilson, K., Yoneyama, K., Boyer, A., & Steffen, H. (2002). The haptic Mller-Lyer illusion insighted and blind people. Perception, 31(10), 1263–1274. doi: 10.1068/p3340

Simonyan, K., & Zisserman, A. (2014, September). Very Deep Convolutional Networks for Large-Scale Image Recognition. 1409.1556 [cs]. Retrieved 2019-06-07, from http://arxiv.org/abs/1409.1556 (arXiv: 1409.1556)

Szegedy, C., Vanhoucke, V., Ioffe, S., Shlens, J., & Wojna, Z. (2015, December). Rethinking the Inception Architecture for Computer Vision. 1512.00567 [cs]. Retrieved 2019-06-07, from http://arxiv.org/abs/1512.00567 (arXiv: 1512.00567)

Yamins, D. L. K., Hong, H., Cadieu, C. F., Solomon, E. A., Seibert, D., & DiCarlo, J. J. (2014, June). Performance-optimized hierarchical models predict neural responses in higher visual cortex. Proceedings of the National Academy of Sciences, 111(23), 8619–8624. Retrieved 2019-06-08, from https://www.pnas.org/content/111/23/8619 doi:10.1073/pnas.1403112111

